# The spectral features of EEG responses to transcranial magnetic stimulation of the primary motor cortex depend on the amplitude of the motor evoked potentials

**DOI:** 10.1101/133769

**Authors:** Matteo Fecchio, Andrea Pigorini, Angela Comanducci, Simone Sarasso, Silvia Casarotto, Isabella Premoli, Chiara-Camilla Derchi, Alice Mazza, Simone Russo, Federico Resta, Fabio Ferrarelli, Maurizio Mariotti, Ulf Ziemann, Marcello Massimini, Mario Rosanova

**Author notes:** These authors contributed equally to this work (Matteo Fecchio, Andrea Pigorini). CORRESPONDING AUTHOR: Mario Rosanova, Department of Biomedical and Clinical Sciences “L. Sacco”, University of Milan, via GB Grassi 74, 20157 Milan, Italy.

## Abstract

Transcranial magnetic stimulation (TMS) of the primary motor cortex (M1) can excite both cortico-cortical and cortico-spinal axons resulting in TMS-evoked potentials (TEPs) and motor-evoked potentials (MEPs), respectively. Despite this remarkable difference with other cortical areas, the influence of motor output and its amplitude on TEPs is largely unknown. Here we studied TEPs resulting from M1 stimulation and assessed whether their waveform and spectral features depend on the MEP amplitude. To this aim, we performed two separate experiments. In experiment 1, single-pulse TMS was applied at the same supra-threshold intensity on primary motor, prefrontal, premotor and parietal cortices and the corresponding TEPs were compared by means of local mean field power and time-frequency spectral analysis. In experiment 2 we stimulated M1 at resting motor threshold in order to elicit MEPs characterized by a wide range of amplitudes. TEPs computed from high-MEP and low-MEP trials were then compared using the same methods applied in experiment 1. In line with previous studies, TMS of M1 produced larger TEPs compared to other cortical stimulations. Notably, we found that only TEPs produced by M1 stimulation were accompanied by a late (∼300 ms after TMS) event-related desynchronization (ERD), whose magnitude was strongly dependent on the amplitude of MEPs. Overall, these results suggest that M1 produces peculiar responses to TMS possibly reflecting specific anatomo-functional properties, such as the re-entry of proprioceptive feedback associated with target muscle activation.

## 1. INTRODUCTION

The combination of Transcranial Magnetic Stimulation with electroencephalogram (TMS/EEG) allows recording the immediate response of the cerebral cortex to a focal perturbation with good spatial and temporal resolution [1]. Indeed, TMS-evoked EEG potentials (TEPs) provide a reliable read-out of the reactivity of cortical circuits provided that they are not confounded by scalp muscle artifacts or spurious sensory activations [2–4]. Typically, TEPs can last for hundreds of milliseconds and are characterized by sustained increases of power in frequency bands that specifically depend on the cortical target [5–8].

Over the last decade, separate TMS/EEG studies have been conducted both on primary motor and sensory cortical areas [5,9–17] as well as associative cortical areas [6,7,18–22]. In this respect, the primary motor cortex (M1) has been largely used as the elective experimental model to study brain reactivity to TMS. However, The EEG response of M1 to TMS may represent a very special, instead of a representative case. In fact, the stimulation of M1 above resting motor threshold (RMT) [23,24] implies not only the excitation of cortico-cortical or cortico-thalamic circuits [5,9,13], but also the excitation of the corticospinal tract [25], which may, in turn, elicit motor evoked potentials (MEPs) and somatosensory feedback [26,27]. Moreover, M1 presents unique features in term of cytoarchitectonics, due to the strong presence of large V layer pyramidal cells [28,29] and the synaptic density/efficacy of corticospinal connections [30]. Previous studies have demonstrated that power and phase of ongoing EEG oscillations can modulate the amplitude of MEPs [14,31–33]. However, so far nobody investigated if and how the amplitude of MEPs influences the waveform and the frequency content of TEPs.

The present investigation aims at evaluating the specific characteristics of M1 TEPs compared to those of other cortical regions that do not elicit any peripheral output. Specifically, we performed two different experiments in order (i) to compare M1 TEPs to those elicited by prefrontal, premotor and parietal cortex stimulation and (ii) to test whether the TEPs generated by M1 stimulation are influenced by the related MEP amplitude. Compared to the other cortical areas, we found that TEPs recorded after stimulation of M1 were larger and characterized by a significant late broadband reduction of power (event-related desynchronization - ERD). Importantly, we observed that larger MEPs were associated with larger TEPs and deeper ERD.

## 2. MATERIALS AND METHODS

### 2.1. Subjects

Six right-handed healthy subjects (3 female; age: 28 ± 3.6 years, values are given as mean ± standard error here and in the following) took part in experiment 1 and a different group of six right-handed healthy subjects (2 female; age: 32 ± 3.0 years) participated in experiment 2 (see Experimental Procedure) after giving written informed consent. All volunteers were screened for contraindications to TMS during a physical and neurological examination [34]. Exclusion criteria included history of neurological or psychiatric disease, of CNS active drugs and abuse of any drug (including nicotine and alcohol). Experiments were approved by the Ethics Committee Milano Area A.

### 2.2. TMS targeting

In both experiments, a focal figure-of-eight coil (mean/outer winding diameter 50/70 mm, biphasic pulse shape, pulse length 280 µs, focal area of the stimulation 0.68 cm^2^) connected to a Mobile Stimulator Unit (eXimia TMS Stimulator, Nexstim Ltd) was used to deliver single-pulse TMS. Stimulation sites included the middle frontal gyrus (Brodmann area [BA] 46), the superior frontal gyrus (BA 6), the precentral gyrus (BA 4) for M1, and the superior parietal gyrus (BA 7) on the left (motor dominant) hemisphere. All of these areas were anatomically identified on a T1-weighted individual magnetic resonance brain scan acquired with an Ingenia 1.5 T (Philips) scanner. For the left motor area, we stimulated the hand area corresponding to the abductor pollicis brevis muscle (APB) of the right hand, which was determined as the site where TMS consistently produced a selective muscle twitch. Stimulation parameters were controlled by means of a Navigated Brain Stimulation (NBS) system (Nexstim Ltd.) that employed a 3D, frameless infrared tracking-position sensor unit to display online, on the individual MRI scan, the position of the TMS coil with respect to the subject’s head. NBS also estimated online the distribution and intensity (expressed in V/m) of the intracranial electric field induced by TMS and allowed to reliably control the stimulation coordinates, within and across sessions [35,36], by signaling in real-time any deviation from the designated target (error <3 mm). In order to standardize stimulation parameters, the maximum electric field was always kept on the convexity of the targeted gyrus with the direction of the induced current perpendicular to its main axis.

### 2.3. TEP and MEP recording

TEPs were recorded with a 60-channel TMS-compatible amplifier (Nexstim Ltd.), that prevents amplifier saturation and reduces, or abolishes, the magnetic artefacts induced by the coil’s discharge [37]. The EEG signals were bandpass-filtered 0.1-350 Hz, sampled at 1450 Hz and referenced to an additional forehead electrode. Horizontal and vertical eye movements were recorded using two additional electrooculogram (EOG) sensors. Impedances at all electrodes were kept < 5 kΩ. As in previous studies, a masking noise capturing the specific time-varying frequency components of the TMS click was played via earphones throughout the entire TMS/EEG sessions to avoid contamination of the EEG signal by auditory potentials evoked [6,18,36].Moreover, bone conduction was attenuated by placing a thin layer of foam between coil and scalp [38].

A 6-channel eXimia electromyography (EMG) system (3000 Hz sampling rate and 500 Hz cutoff for low-pass filtering) was used to record MEPs. Ag-AgCl self-adhesive electrodes were placed over the right APB muscle according to the belly–tendon montage [39].

### 2.4. Experimental procedures

During all TMS/EEG recordings, subjects were seated on a comfortable reclining chair, with eyes open and with the right hand positioned on a pillow placed over their lap. During stimulation of M1, subjects were instructed to keep the target muscle relaxed while EMG was continuously monitored on a computer screen. In experiment 1, for every subject we performed 4 TMS/EEG measurements in which 4 cortical target were stimulated with a random order across subjects. For each TMS/EEG measurement we collected ∼250 trials at an estimated electric field on the cortical surface of 120 V/m. This intensity range is considered effective to produce significant EEG responses [35,36] and, at the same time, always evoked a clear but selective APB muscle twitch when targeting M1 (i.e. supra-threshold intensity, where the resting motor threshold [RMT] is defined as the minimum intensity necessary to elicit a peak-to-peak MEP amplitude higher than 50 µV in at least 5 out of 10 subsequent trials while muscle target is at rest, as assessed by the relative frequency method [23,24]). For each stimulation target, 250 TMS pulses were delivered at an inter-stimulus interval (ISI) randomly jittered between 3000 and 3300 ms. In experiment 2, M1 was stimulated with an ISI of 5000-5300ms (random jittering) to conform to the typical ISI used in the literature of M1 stimulation [40,41]. In experiment 2, a total number of 500 TMS pulses were delivered at RMT, corresponding to a mean estimated electric field of 93 ± 5.9 V/m across subjects.

### 2.5. Data analysis

For both experiments, data analysis was performed using Matlab R2012a (The MathWorks). Artifact-contaminated channels and trials were manually rejected by visual inspection [22] (in experiment 2, trial rejection also involved the visual inspection of single-trial MEPs). Then, EEG data were band-pass filtered (1-80 Hz, Butterworth, 3rd order), half-sampled at 725 Hz and segmented in a time window of ± 600 ms around TMS pulses. Bad channels were interpolated using EEGLAB spherical interpolation function [42] and signals were average re-referenced and baseline corrected. Independent component analysis (ICA) was applied in order to remove residual eye blinks/movements and scalp muscle activations. In experiment 1, TEPs were obtained by averaging a minimum of 100 artifact-free single trials (170 ± 11 for prefrontal stimulation, 158 ± 16 for premotor stimulation, 118 ± 15 for M1 stimulation, 139 ± 18 for parietal stimulation, respectively).

In experiment 2, EMG traces were filtered (2 Hz high-pass, Butterworth, 3rd order) and segmented in a time window of ± 150 ms around the TMS pulse. Based on their peak-to-peak amplitude distribution across a large number of artifact-free trials (396 ± 19), for each session we averaged separately the 100 trials with the largest MEP amplitude (high-MEP condition) and the 100 trials with the smallest MEP amplitude (low-MEP condition; an example is shown in Fig 2A).

In order to assess and compare the local TMS-evoked activity between stimulation sites and between high-MEP and low-MEP conditions, we calculated the Local Mean Field Power (LMFP) computed as the square root of squared TEPs averaged across the four channels located under the stimulation coil (similar to [43]) and pertaining to the area of the scalp over each of the four cortical targets (prefrontal, premotor, motor and parietal).

Bootstrap statistics with *p* < 0.01 (number of permutations = 1000) were applied on LMFP time-series. Only the LMFP values significantly different from the baseline (−500 to −100 ms) were averaged between +20 and +350 ms and used in the analysis measuring the local TEPs amplitude.

Spectral features were evaluated by computing the event-related spectral perturbation (ERSP) [42] between 8 and 45 Hz [6–8] after time-frequency decomposition using Wavelet transform (Morlet, 3.5 cycles). Absolute spectra normalization was applied first at the single-trial level performing a full-epoch length single-trial correction [44] and then by a pre-stimulus baseline correction (−500 to −100 ms) on the resulting ERSP averaged across trials. Finally, similar to LMFP, only significant ERSP values surviving bootstrap-based statistics (α < 0.05, number of permutations = 500) with respect to baseline were considered in the analysis. Specifically, for each of the simulated cortical areas, we averaged the ERSP in the 8-45 Hz frequency range and in the time window between +200 and +350 ms across the same four channels selected for LMFP calculation.

### 2.6. Statistics

In experiment 1 significant differences among areas were assessed, for both LMFP and ERSP, by means of one-way analysis of variance (ANOVA) and pairwise post-hoc two-tailed t-tests were used (Bonferroni corrected). In experiment 2, statistical comparison between high-MEP and low-MEP conditions was performed on the LMFP and ERSP by means of paired t-test.

## 3. RESULTS

### 3.1. Experiment 1: M1 response to TMS is larger and characterized by distinct spectral features compared to the prefrontal, premotor, and parietal cortex responses.

The local amplitude of TEPs, as measured by the significant LMFP values averaged between +20 and +350 ms, was significantly different among areas (F_(3,20)_ = 18.9, p = 4.8 *10^-6^). Specifically, M1 stimulation elicited larger TEPs compared to each of the other stimulated sites (p < 0.01, Bonferroni corrected), while the LMFP comparison of prefrontal, premotor and parietal targeting did not show any significant differences (Fig 1C). The same results were obtained by contrasting the TEPs global amplitude across all 60 channels elicited by each stimulation site by means of the corresponding global mean field power (GMFP; see Fig S1).

**Fig 1.**
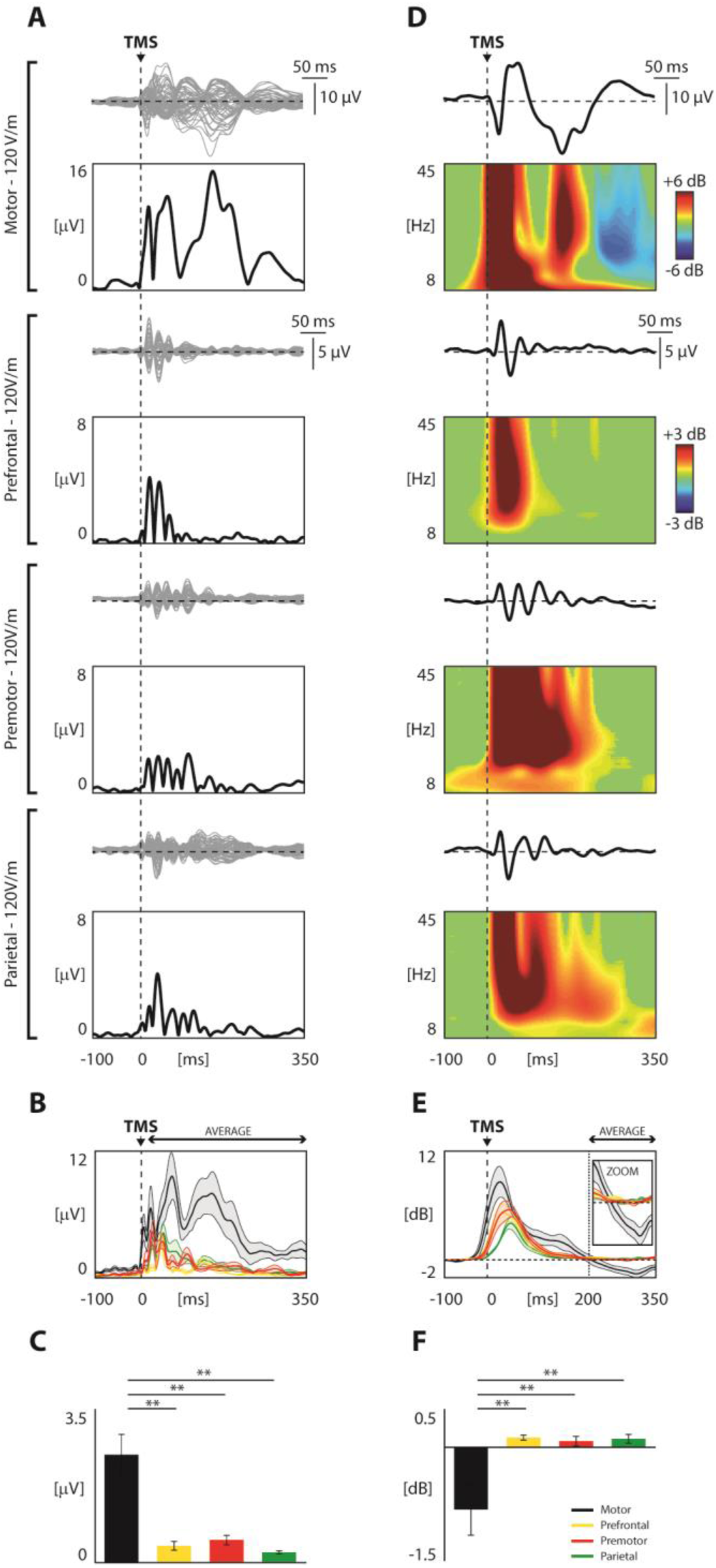
Comparison of TMS-evoked EEG potentials recorded from different cortical areas employing Local Mean Field Power (LMFP) and Event-related Spectral Perturbation (ERSP). **Panel A**. For each stimulated area, TMS-evoked EEG responses are shown from one representative subject. Butterfly plots of all channels are displayed (top panels, grey traces), together with the corresponding LMFP (bottom panels, black traces). The dashed vertical line indicates the timing of the TMS pulse. **Panel B**. Grand-average of LMFP for each stimulated area. Thick traces indicate the grand-average LMFP across subjects (±SE, color-coded shaded regions). Responses recorded after the stimulation of different cortical areas are color coded as follows: motor in black, prefrontal in yellow, premotor in red, parietal in green. **Panel C**. For each stimulated area, the LMFP values averaged between 20 and 350 ms post-TMS are shown in the bar histogram (mean ± SE). Asterisks indicate statistically significant differences (** p<0.01, pairwise comparison, Bonferroni corrected). Bars are color coded as in Panel B. **Panel D**. Black traces represent TMS-evoked EEG responses of a representative subject (same as in panel A) for one of the four channels closest to the stimulation site, with the corresponding ERSP shown below. A Wavelet Transform (Morlet, 3.5 cycles) has been applied at the single trial level. Significance threshold for bootstrap statistics is set at α<0.05. Non-significant activity is set to zero (green), red colors indicate a significant increase with respect to the baseline, while blue colors indicate a significant reduction compared to the baseline. As in Panel A, the dashed vertical line indicates the time of the TMS pulse. **Panel E**. The averaged ERSP of the four channels located under the stimulation coil (between 8 and 45 Hz) is presented for each stimulated area (color coding as in panel B and C). Each thick line indicates the grand-average across subjects (± SE, color-coded shaded regions). The same traces are enlarged in the inset (time scale from 150 to 350 ms; power scale from −2 to 2 dB). The dashed vertical line indicates the average time in which the ERD occurs. **Panel F**. Using the same color coding as in panel B, C and E, bars indicate, for each stimulated area, the grand-average (±SE) of the averaged ERD (ERSP between 200 and 350 ms post-TMS). Asterisks indicate statistically significant differences (** p<0.01, pairwise comparison, Bonferroni corrected).

With respect to the EEG responses in the time-frequency domain (ERSP) we observed that all targeted cortical areas responded to TMS with a broadband increase of spectral power lasting up to ∼200 ms (Fig 1E). After this first activation, spectral power returned to baseline in all targeted cortical areas except for M1, which showed a statistically significant event related desynchronization (ERD; blue color in top panel of Fig 1D). Across subjects, this ERD reached maximum values at about 300 ms post-TMS (310 ± 3.5 ms), as shown by the grand-average ERSP cumulated over the entire 8-45 Hz frequency range (Fig 1E). Indeed, statistical analysis showed that the modulation of spectral power was significantly different among areas (F_(3,20)_ = 7.06, p = 0.002) when averaged between +200 and +350 ms. Specifically, M1 modulation of power was significantly different from all the other stimulated sites, while the contrasts between prefrontal, premotor and parietal targets did not show any significant differences (Fig 1F).

### 3.2. Experiment 2: M1 responses to TMS are influenced by MEP amplitude.

First, we replicated the M1-related ERD observations found in experiment 1 using a longer ISI (5000-5300 ms). Specifically, we observed that stimulating every 5000-5300 ms, M1 TEPs were characterized by a statistically significant late ERD averaged across the four channels closest to the stimulation site in the range of 8-45 Hz similar to that obtained using a shorter ISI (3000-3300 ms; Fig S2).

Most importantly, for each session, the high number of recorded trials (n = 500) allowed to extract and analyze separately two subsets of 100 trials each, selected on the basis of the largest (high-MEP) and the smallest (low-MEP) single-trial MEP amplitude respectively. Across subjects, in the high-MEP condition the lowest MEP amplitude was on average 504 µV (± 174 µV), while in the low-MEP condition the highest amplitude was on average 165 µV (±38 µV), therefore ensuring the absence of any overlap between high and low MEP conditions (Fig 2A, left panel). The average MEP amplitude for the high-MEP and low-MEP conditions were 1026 µV (range 223-1527 µV) and 29 µV (range 4-72 µV), respectively (Fig 2D, left panel). The overall amplitude of TEPs, as measured by the average of significant LMFP values between +20 and +350 ms, was significantly reduced (20.6% ± 2.2; Fig 2C) in low-MEP condition as compared to high-MEP condition (one-tail paired t-test, p<0.01) at the group level (Fig 2D, middle panel).

Regarding the effects of MEP amplitude on the spectral features of the local M1 EEG response, we found that the amount of ERD was significantly reduced (72.4% ± 18.3; Fig 2D, right panel) in the low-MEP condition as compared to the high-MEP condition (one-tail paired t-test, p<0.01). Notably, a topographic statistical analysis (Fig 3) indicated that the significant ERD reduction was confined (one-tail paired t-test, p<0.01) to the channels overlying the left motor area (C3, C5, FC3, FC5, and FC1).

**Fig 2.**
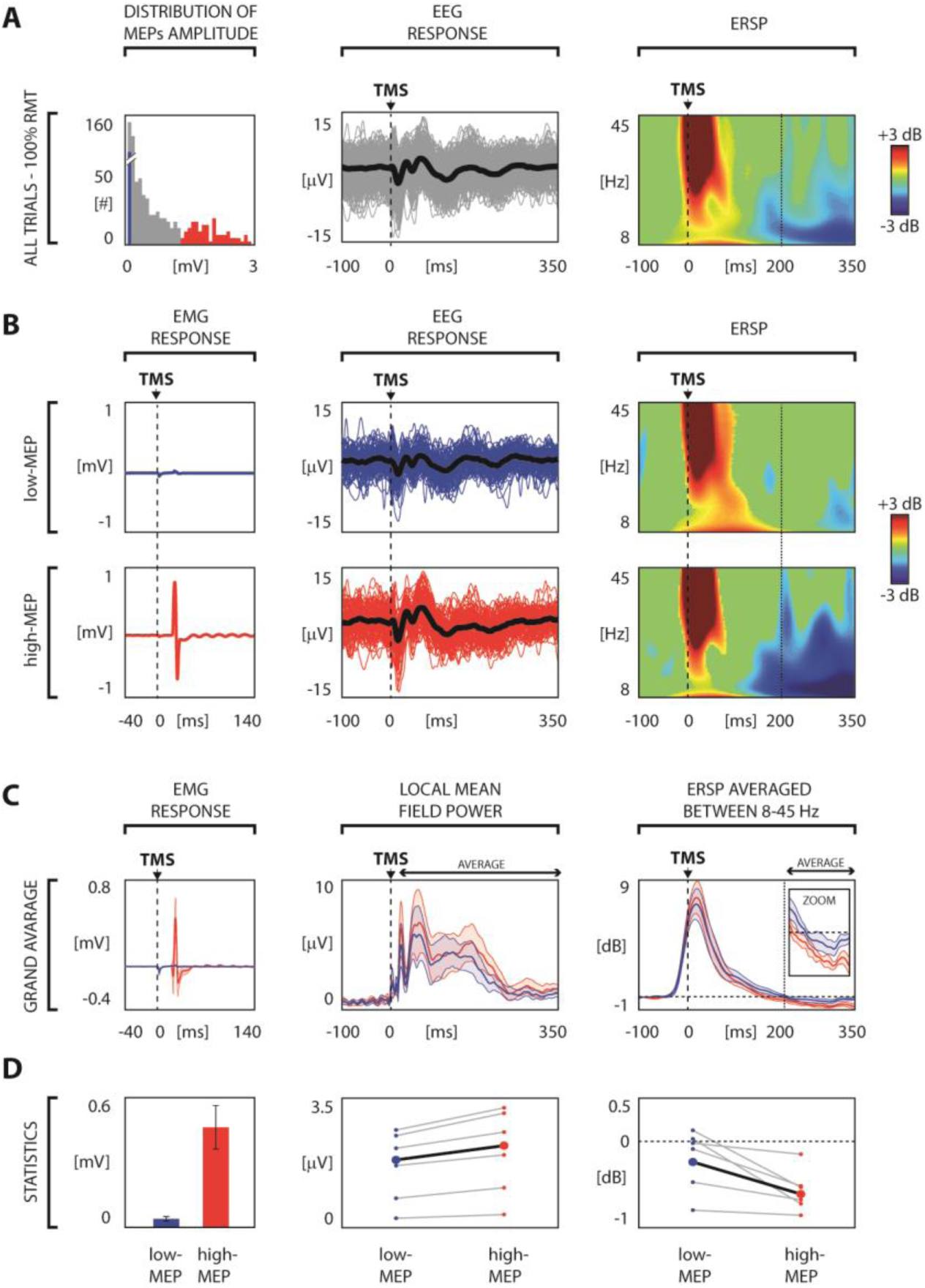
TMS-evoked EEG responses over left M1: comparison between high-MEP and low-MEP conditions. Panel A. Left panel shows the distribution of the peak-to-peak MEPs amplitude of the all artifact-free trials for one representative subject. Blue bars correspond to 100 trials in which TMS elicited the smallest APB motor responses (low-MEP) and red bars corresponds to 100 trials in which TMS generated the largest APB motor responses (high-MEP). Then, from left to right, the EEG single trials (thin lines) and the average response (thick line) of the channel closest to the stimulation site (C3 scalp derivation) and the corresponding ERSP are shown. **Panel B.** For the same representative subject of Panel A, low-MEP (top panel) and high-MEP conditions (bottom panel) are compared. From left to right, the average MEP, the EEG single trials (thin lines) and average TEPs (thick line) recorded from the electrode closest to the stimulation site (C3 scalp derivation) and the corresponding ERSPs are shown. **Panel C.** MEP, LMFP and averaged ERSP derived from low-MEP (blue) and high-MEP (red) trials are compared. Each thick line indicates the grand-average across subjects (±SE, color-coded shaded regions). The averaged ERSP traces are enlarged in the inset (time scale from 150 to 350 ms; power scale from −1 to 1 dB). **Panel D.** From left to right: average (±SE) across subjects of the MEP peak-to-peak amplitude, individual averaged LMFP between 20 and 350 ms and the individual averaged ERSP between 200 and 350 ms are presented. Small circles and grey lines indicate single subject values, while large circles and black lines indicate grand-average values across subjects. Statistical analysis by means of paired t-test resulted in significant differences for MEP (p<0.05), LMFP (p<0.01) and ERSP (p<0.01).

**Fig 3.**
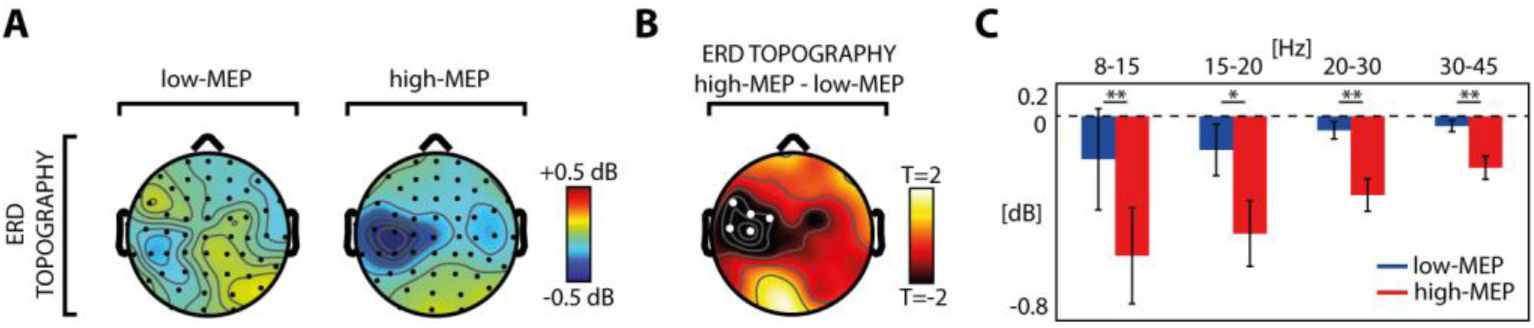
Comparison between low-MEP and high-MEP conditions across channels and frequency bands. Panel A. The broadband (8-45 Hz) ERD topography (ERSP averaged between 200 and 350 ms) of the grand-average across subjects derived from both low-MEP (left) and high-MEP conditions (right) is shown. **Panel B** displays the topographic distribution of the t-values from a paired t-test (p<0.05) together with the statistical differences between the broadband ERD in the two conditions. The statistically significant electrodes (white dots) indicate that this broadband reduction was confined over the left motor area (C3, C5, Fc3, Fc5, and Fc1 scalp derivations). **Panel C.** For the significant channels of Panel B, the ERSP averaged between 200 and 350 ms across channels in the low-MEP (blue) and high-MEP (red) conditions over four EEG frequency bands (8-15Hz, 15-20Hz, 20-30Hz, 30-45 Hz) are shown in the bar histogram (±SE). Asterisks indicate statistically significant differences (* p<0.05, ** p<0.01, paired t-tests).

Finally, we split the average ERD over these five channels into four frequency bands (8-15 Hz, 15-20 Hz, 20-30 Hz, 30-45 Hz) and tested the differences across frequencies between high-MEP and low-MEP conditions using a two-way within subject ANOVA with MEP conditions and frequencies as main factors. Results showed a significant main effect of MEP amplitude (F_(1,3)_ = 13.16, p = 0.0008), with a significantly reduced ERD for low-MEP as compared to high-MEP condition across all frequency bands. Conversely, neither the main effect of frequency (F_(1,3)_ = 0.29, p = 0.83) nor the interaction between frequency and MEP amplitude (F_(1,3)_ = 0.59, p = 0.62) were significant.

## 4. DISCUSSION

In the present study, we investigated the peculiar features of the EEG responses of M1 to TMS and we asked whether those features are related to the amplitude of MEPs. To this aim we performed two series of experiments: in experiment 1 we compared TEPs of M1 with those of other cortical areas (premotor, prefrontal and parietal) in the time-frequency domain. Here stimulations were performed at an intensity of 120 V/m, which is largely above the threshold to reliably elicit a MEP. In experiment 2, we stimulated M1 at RMT and compared TEPs acquired within the same session and classified as low-MEP and high-MEP based on MEP amplitude. We found that: (i) according to previous studies [45] M1 TEP amplitude is larger as compared to any of the other stimulated areas, (ii) only the M1 response is associated with a late ERD (∼300 ms), which is clearly modulated by the amplitude of MEPs.

Behavioral and electrophysiological measurements suggested that M1 is more excitable than other cortical areas [45–47]. For instance, previous works showed that TMS delivered above RMT over M1 and prefrontal cortex resulted in larger global response (GMFP) for the stimulation of M1 [45,47,48]. Here we confirmed and extended these results (experiment 1) by comparing TEPs of M1 to those generated by stimulating the premotor and posterior parietal cortices, which are usual targets in TMS/EEG experiments [6–8,49]. We observed that the M1 LMFP and GMFP to supra-threshold TMS (120 V/m as estimated by the neuronavigation system) are significantly larger than those measured after the stimulation of premotor and parietal cortices (Fig 1A-C, and Fig S1).

Most important, the analysis of TEPs in the time-frequency domain showed that only the stimulation of M1 is associated to a statistically significant late (∼300ms) broadband ERD (Fig 1D-F), whose scalp topography is confined to the EEG electrodes overlying the stimulated sensory-motor areas (Fig 3). In terms of spectral and topographical features, the ERD we observed closely resemble the localized desynchronization of the ongoing EEG oscillations in the low and high µ-bands induced by somatosensory stimulation [50] and by the execution of a voluntary movement [51]. Along the same lines, other studies showed that ERD in the β-band (15-30 Hz) recorded from sensory-motor cortices is associated with electrical nerve stimulation [52] and mechanical finger stimulation [53], as well as with movement and motor imagery [54,55].

Thus, the ERD that hallmarks the electrical M1 response we recorded after supra-threshold TMS may be strongly contributed by the activation of specific cortico-spinal circuits [56] and the sensory feedback from the activated muscle [50,53,57]. In order to further test this hypothesis, we performed a second set of measurements (experiment 2) in which TEPs following the stimulation of M1 at an intensity corresponding to RMT and were ranked based on the occurrence of high amplitude and low amplitude MEPs. Results clearly showed that in the high-MEP condition (i.e. when the cortico-spinal tract is more activated and the proprioceptive sensory feedback is stronger) the EEG response to TMS was significantly larger with respect to the low-MEP condition (i.e. when cortico-spinal tract is less activated and the proprioceptive sensory feedback is weaker), both in the early (as in [14]) and in the late TEP components (Fig 2C). Notably, also M1 ERD was influenced by the amplitude of MEPs, thus confirming a relationship between M1 EEG response to TMS and the effect of the stimulation at the peripheral level (Fig 2). Interestingly, when the stimulation of M1 did not trigger any muscular activation (as in [7]), the ERD was absent, confirming that the occurrence of ERD is only present when the stimulation actively involves the cortico-spinal tract (regardless the specific targeted muscle) and is associated with the proprioceptive sensory feedback, and absent otherwise (Fig S3).

More in general, the peculiar spectral properties of M1 response to TMS could be related to specific anatomo-functional features of the motor cortex, such as its high level of connectedness previously demonstrated by studies combining TMS with functional MRI [56,58]. In this case, the larger M1 EEG responses may be influenced by the activation of specific cortico-cortical pathways which involves areas directly connected to M1 such as the primary somatosensory cortex [32], the M1 contralateral to the stimulation, the supplementary motor and premotor areas [13,56,59]. With respect to the timing of this late ERD found in previous studies and confirmed in the present work, its latency is consistent with the time interval required for the elaboration of the subjective awareness of somatosensory stimuli [60] further implying the role of sensory feedback on the specific M1 response to TMS.

In addition to these macro-anatomical aspects, also the peculiar M1 cytoarchitectonics may play a role. Indeed, magnetic stimulation seems to be more effective in exciting longitudinally oriented pyramidal cells which have a large-diameter myelinated axon and a wider dendritic tree [2,61]. Thus, the direct activation of giant Betz cells of layer Vb, whose strong presence is a unique feature of M1 [28,29], may contribute to the distinct features of M1 TEPs. Altogether, these findings confirmed that the M1 EEG responses to TMS show peculiar features strongly related to the concurrent activation of a peripheral output. Future studies should explore the contributions of cytoarchitectonics and structural connectivity to the specificity of TEPs elicited in other cortical areas, such as the occipital cortex, towards the development of a non-invasive perturbational atlas.

## ACKNOWLEDGMENT

We thank Sasha D’Ambrosio, Thierry Nieus and Renate Rutiku for insightful discussion and comments on the manuscript. This study has been partially funded by the EU grant H2020-FETOPEN-2014-2015-RIA n. 686764 “Luminous”, the grant "Sinergia" CRSII3_160803/1 founded by the Swiss National Science Foundation, the James S. McDonnell Foundation Scholar Award 2013, the EU grant H2020 grant agreement 720270-Human Brain Project SGA1 (to M.Mas.), the Grant "Giovani Ricercatori" GR-2011-02352031 from the Italian Ministry of Health (to MR).

## SUPPLEMENTARY RESULTS

**TMS over the medial M1 (leg area) leads to and ERD only in the presence of muscle twitch**

Resting Motor Threshold (RMT) is defined as the minimum stimulus intensity that produces a minimal motor evoked response (about 50 µV in at least 5 out of 10 trials) while the target test muscle is at rest [1]. RMT changes depending on the considered target muscle due to scalp-to-cortex distance [2] and higher stimulation intensity is needed to induce a muscle twitch when perturbing the medial portion of the M1 (e.g. the leg motor area). Along the same line, here we show that, when stimulating medial motor cortex, TMS-induced ERD can be obtained just stimulating at the RMT of the targeted muscle (here the right quadriceps). To do this, in one representative subject, once obtained RMT by stimulating APB area (RMT_APB_), we moved to a more medial area, performing a first TMS/EEG recording by targeting the portion of M1 corresponding to right quadriceps. Then, we increased the stimulation intensity until reaching a selective MEP of the right quadriceps (RMT_LEG_) muscle without involving APB activation.

**Fig S1. Panel A.**
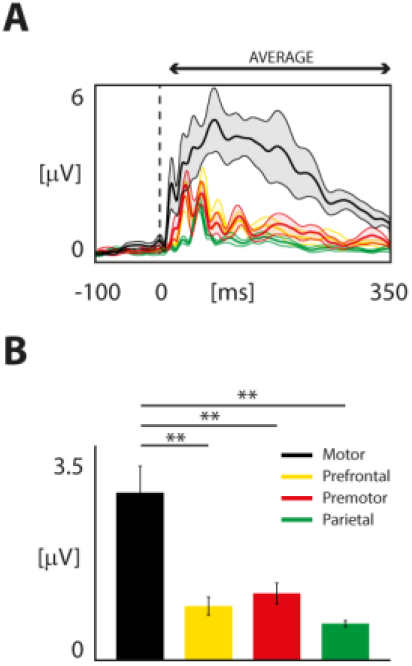
Grand-average of GMFP for each stimulated area. Thick traces indicate the grand-average GMFP across subjects (±SE, color-coded shaded regions). Responses recorded after the stimulation of different cortical areas are color coded as follows: motor in black, prefrontal in yellow, premotor in red, parietal in green. **Panel B**. For each stimulated area, the GMFP values averaged between 20 and 350 ms post-TMS are shown in the bar histogram (mean ± SE). Asterisks indicate statistically significant differences (** p<0.01, pairwise comparison, Bonferroni corrected). Bars are color coded as in Panel A.

**Fig S2.**
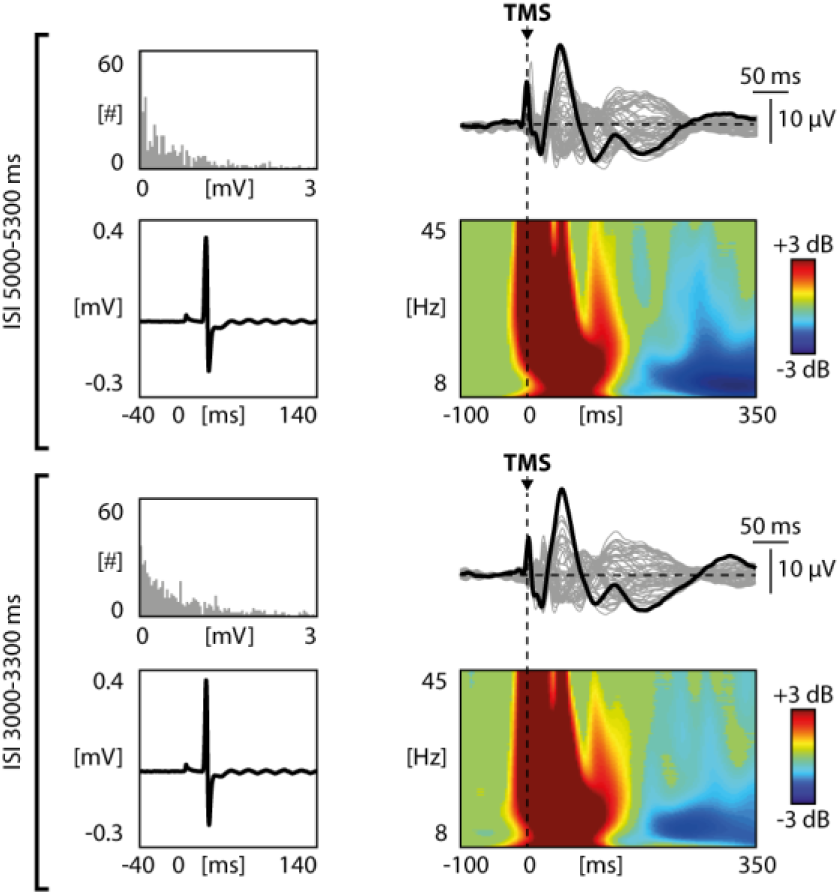
Comparison of 5 second (top panels) and 3 second ISI (bottom panels) in one representative subject. For both ISIs, the distribution of peak-to-peak MEP amplitude of all artifact free trials (left top) and the corresponding average MEP (left bottom), the butterfly plots of all channels (right top, grey traces), the TEPs recorded at the channel closest to the stimulation site (right top, black traces) and the corresponding ERSPs (right bottom) are shown. Wavelet Transform (Morlet, 3.5 cycles) was applied at the single trial level. Significance threshold for bootstrap statistics is set at α<0.05. Non-significant activity was set to zero (green), red colors indicate a significant increase with respect to the baseline, while blue colors indicate a significant reduction with respect to the baseline. The dashed vertical line indicates the timing of the TMS pulse.

**Fig S3.**
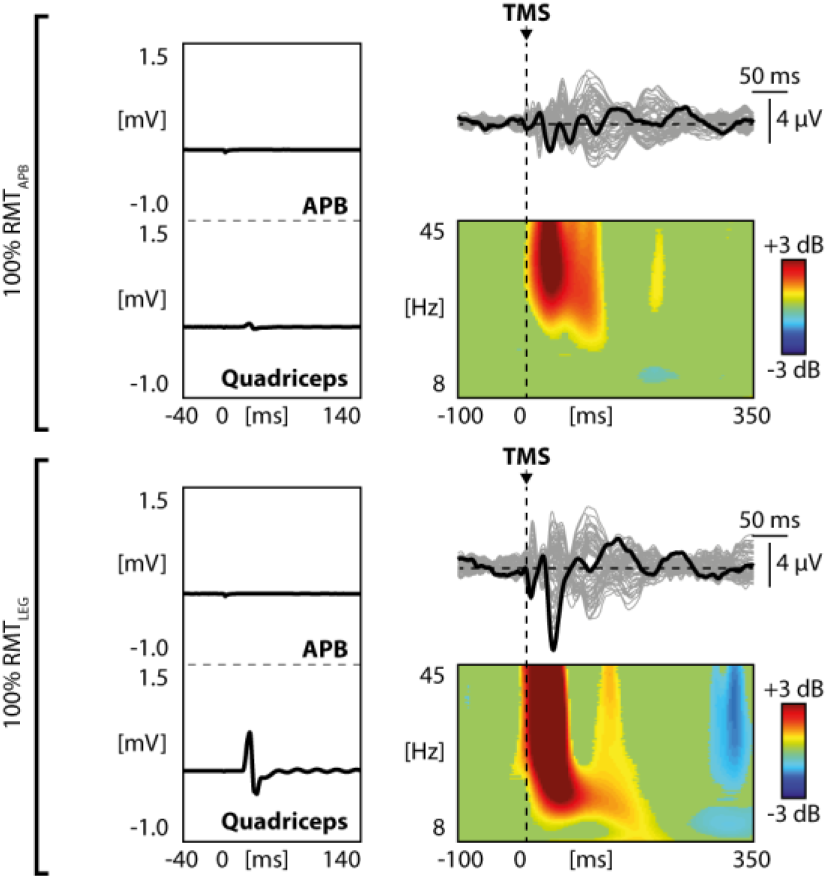
Top and bottom panels show, for one representative subject, the average MEPs of the quadriceps and APB muscles, the butterfly plot of all channels (grey lines), the TEP of the channel closest to the stimulation site (black line) and the corresponding ERSP obtained by stimulating the medial M1 (leg motor cortex) respectively at 100% RMT_APB_ and 100% RMT_LEG_.

